## Supplementary information for "Ultrastructural details of resistance factors of the bacterial spore revealed by in situ cryo-electron tomography"

[a] Univ. Grenoble Alpes, CNRS, CEA, IBS, F-38000 Grenoble, France.
[b] CEITEC-Central European Institute of Technology, Masaryk University, 62500 Brno, Czech Republic.
[c] University of Technology Sydney, 2007 Ultimo, NSW, Australia.
[d] University Grenoble Alpes, CEA, IRIG-MEM, F-38054 Grenoble, France.
[e] School of Life Sciences, University of Warwick, Coventry, UK.
[f] Univ. Grenoble Alpes, CNRS, CEA, EMBL, ISBG, F-38000 Grenoble, France.

**This PDF file includes:**

Supporting text
Figures S1 to S3
Tables S1 to S3
Legends for Movies S1 to S4
SI References

**Other supporting materials for this manuscript include the following:**

Movies S1 to S4

### SUPPORTING INFORMATION TEXT

##### Cryo-FIBM of sporulating *B. subtilis* cells for cryo-electron tomography

Since cryo-FIBM/ET still remains an emerging technique, we have not found any publication that would thoroughly describe preparation of *B. subtilis* samples for cryo-FIBM. We thus tested several parameters to obtain reproducible, thin and robust lamellae of sporulating *B. subtilis* cells. The quality of the prepared lamellae was mostly influenced by the cell concentration, blotting conditions to obtain a uniform monolayer of cells, the vitrification procedure and the milling protocol. Low cell concentration (below $OD_{600nm} \sim 5$) resulted in isolated cells sparsely distributed on the EM grid, while high cell concentration ($OD_{600nm} \sim 20$) allowed an even distribution of cells on the grid surface. To obtain a 5 to 10 µm-thick uniform layer of cells (Fig. S1B), we blotted the back side of the grid with filter paper, and the cell side with a non-absorbent, granular, home-made plastic pad. Importantly, blotting with this device, which exhibited irregularities at its surface, preserved the integrity of the grid better than using smooth plastic polymers like Teflon® pads. The ethane phase had also a notable impact on the stability of the lamellae. We observed that plunge freezing performed at the limit of solidification (milky appearance of ethane), rather than in full-liquid phase (transparent ethane), favored proper sample vitrification. To relieve the tension on the lamellae during and after cryo-FIBM, we milled two trenches on both sides of the lamella (Fig. S1C and S1D) (Wolff et al., 2019). Finally, the last step of polishing was crucial to remove redeposited material from the milling of neighboring lamellae (Moravcová et al., 2021). The best lamellae were obtained using the protocol described in the *Materials and Methods* section. The lamellae thickness was in the range of 150 to 200 nm, as estimated from reconstructed tomograms (Fig. S1E).

### SUPPORTING INFORMATION FIGURES

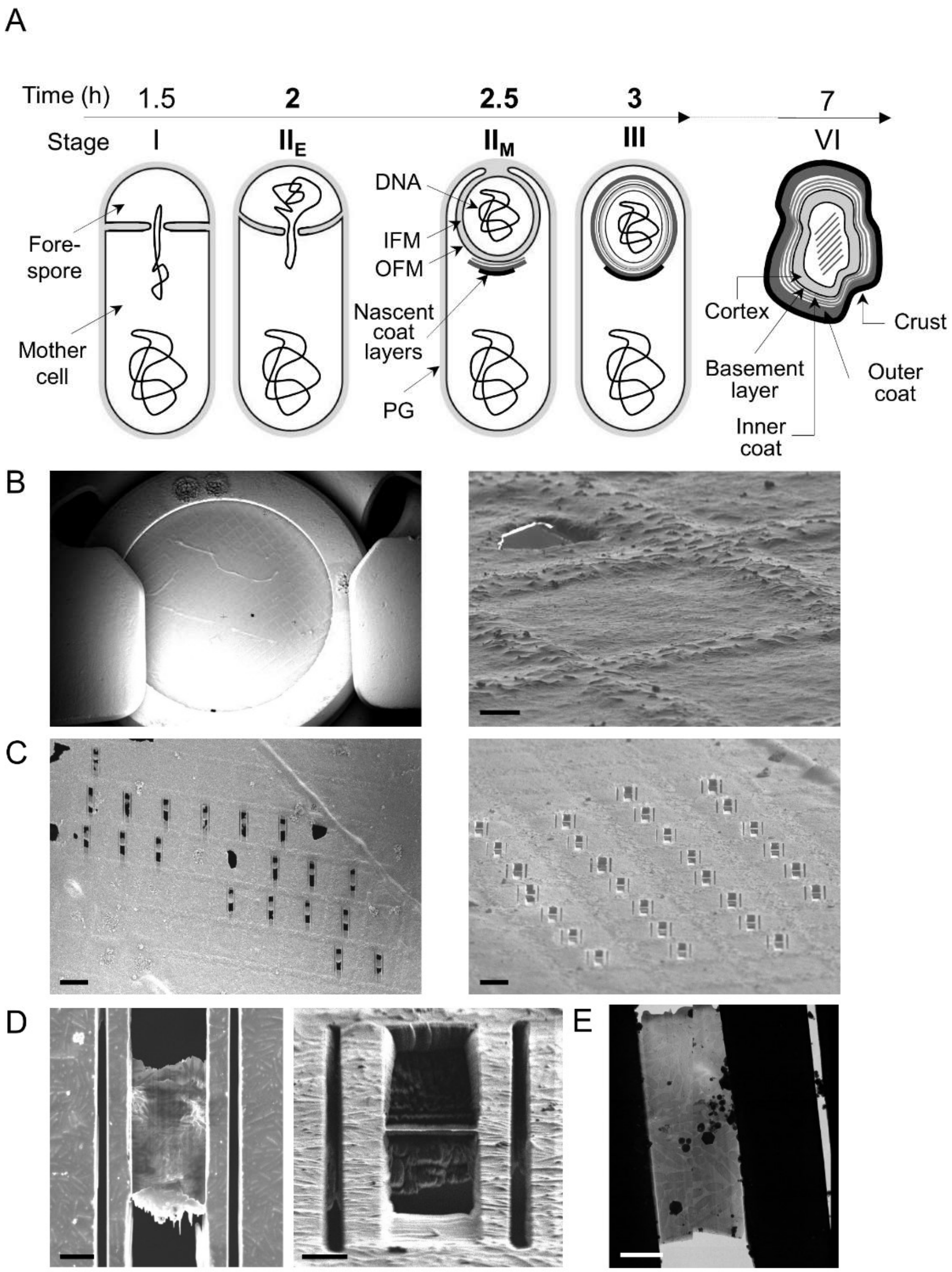


**Figure S1. A.** Schematic illustration of main ultrastructural changes occurring during the different stages of sporulation. IFM, inner forespore membrane; OFM, outer forespore membrane; PG, peptidoglycan. **B.** Full (left panel) and close (right panel, scale bar 5 µm) views of a grid covered with a homogenous layer of *B. subtilis* cells, observed by SEM and FIB, respectively. **C.** Lamellae of vitrified cells milled by FIB, observed from the SEM (left panel) and FIB (right panel) perspective. Scale bars, 50 µm. **D.** Zoom on a FIBM lamella observed from the SEM (left panel) and FIB (right panel) perspective. Scale bars, 5 µm. **E.** TEM view of a FIBM lamella containing vitrified *B. subtilis* cells. Scale bar, 5 µm.

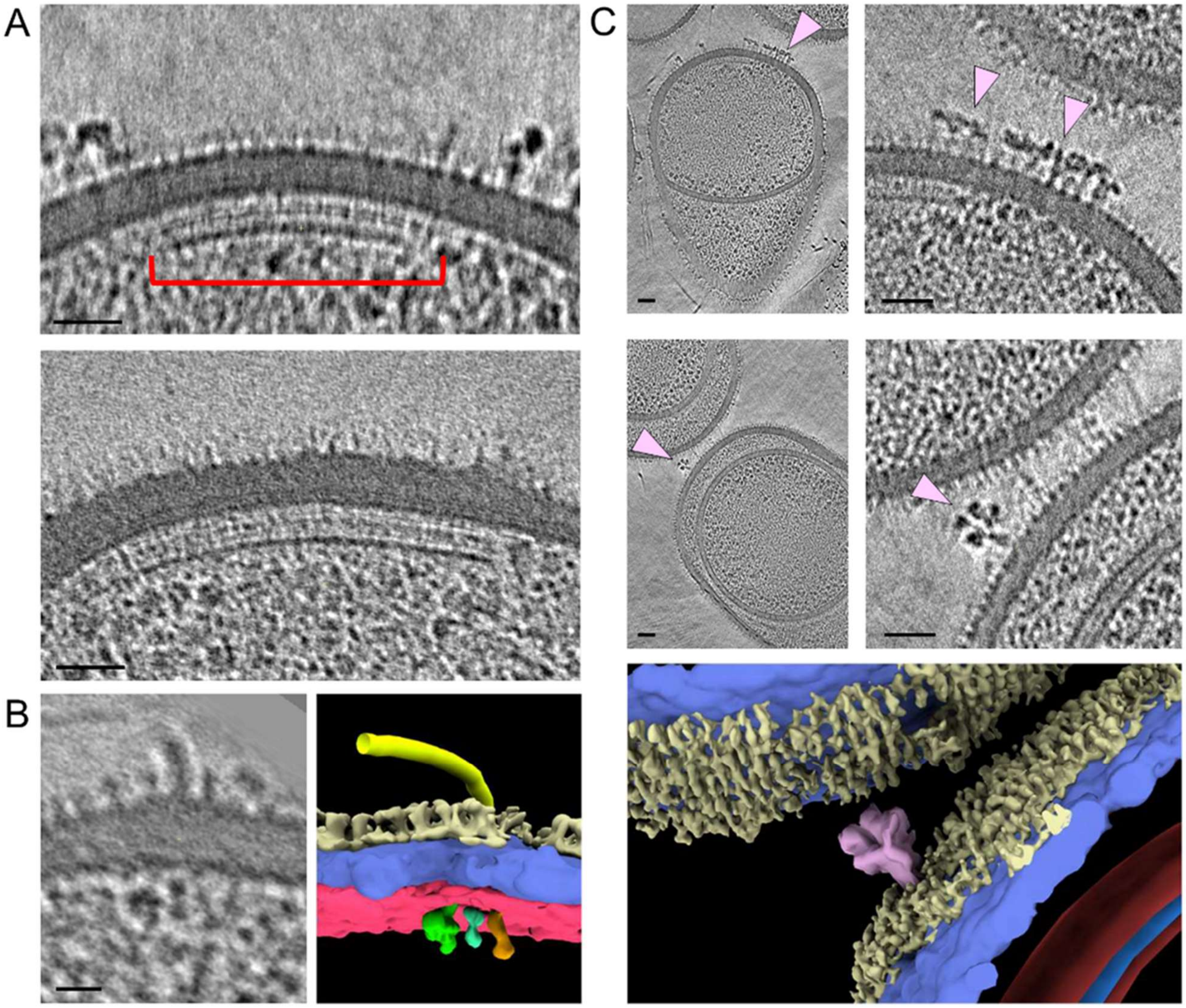


**Figure S2.** Envelope-associated objects revealed by cryo-FIBM/ET. Magnified views of tomogram slices showing chemotaxis arrays (**A**, red staple, scale bars 50 nm), a flagellum (**B**, scale bars 20 nm) and unidentified surface elements (**C**, pink arrowheads, scale bars 50 nm). Segmentation is shown for the surface components. The mother cell membrane is colored in magenta, the PG layer in blue staple and surface glycans in light yellow.

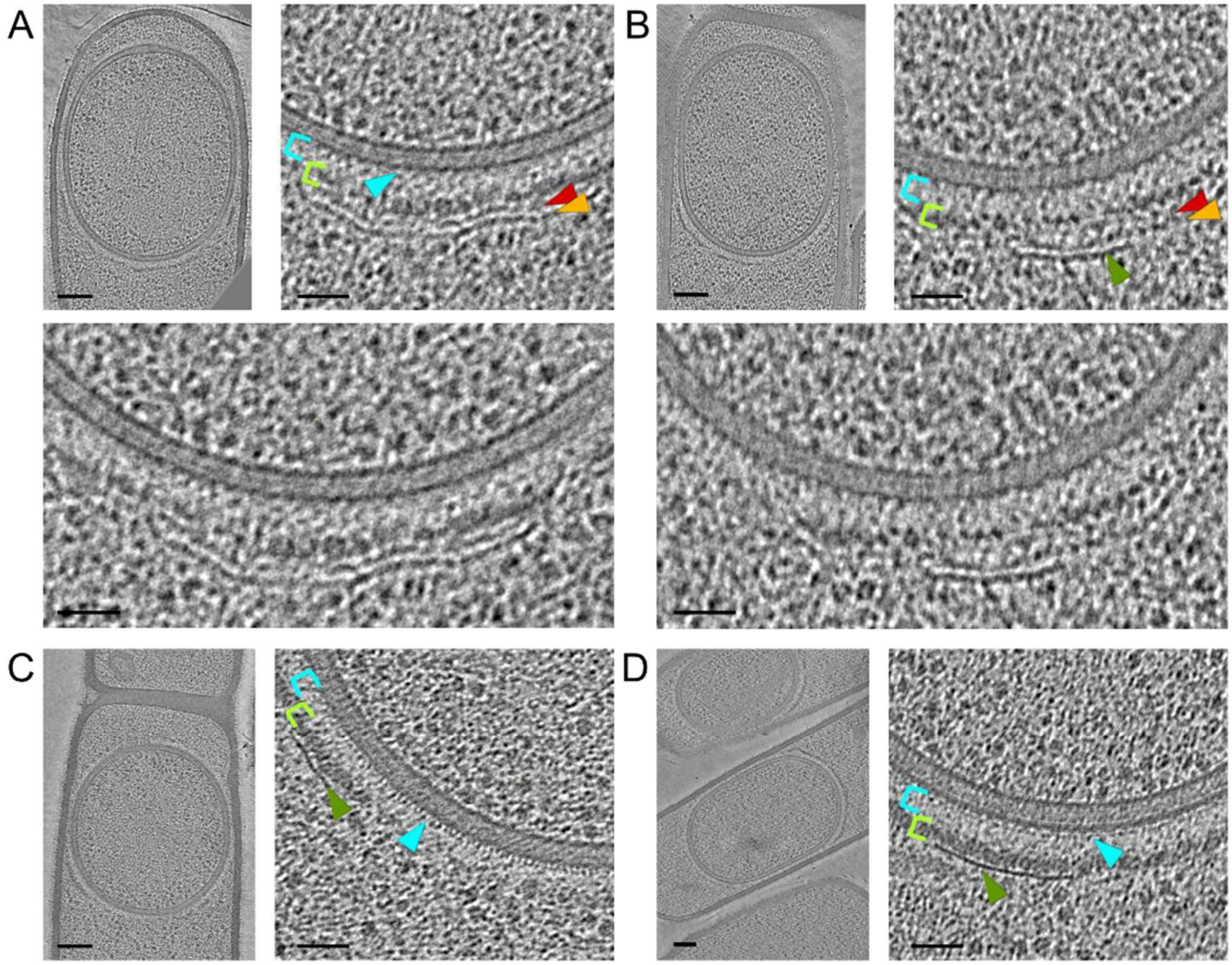


**Figure S3.** Slices through cryo-electron tomograms of *ΔspoIVB* (**A-B**) and *ΔcotE* (**C-D**) sporangia, shown in full view (scale bars, 100 nm) or as magnified views of specific regions (scale bars, 50 nm). Ultrastructures corresponding to the crenelated layer (cyan arrowhead), the light matrix (cyan staple), the dark matrix (lemon staple), the two bead-like layers (red and orange arrowheads) and the dark smooth layer (green arrowhead) are indicated.

**Table S1. Strains used in this study.**

| Construct | Genotype | Source |
|---|---|---|
| ***B. subtilis* strains** | | |
| BDR2413 | 168 | (Zeigler et al., 2008) |
| BCR1661 | 168, *spoIVB::spec* | This study |
| BCR1662 | 168, *spoIVB::spec spoIIQ::kan* | This study |
| BHC77 | 168, *cotE::kan* | This study |
| BBK28 | 168, *spoVID::kan* | (Luhur et al., 2020) |
| BBK33 | 168, *safA::kan* | (Luhur et al., 2020) |
| BJL59 | 168, *spoIVA::cat* | (Luhur et al., 2020) |

**Table S2. Dimensions of cell wall and coat layers measured from cryo-electron tomograms.**

| | **Mean thickness** (nm ± SD)[a] (nb of measurements) | | | | | |
|---|---|---|---|---|---|---|
| **Strain** | **CW** | **CL** | **LM** | **DM** | **BLL** | **DSL** |
| *ΔspoIVB* | 26.9 ± 1.3 (36) | 12.4 ± 0.7 (18) | 31.3 ± 2.4 (24) | 17.8 ± 4.9 (24) | 8.3 ± 0.8 (24) | 7.9 ± 1.0 (24) |
| *ΔcotE*[c] | 29.9 ± 2.4** (30) | 11.2 ± 0.6** (30) | 27.4 ± 2.7** (30) | 19.7 ± 4.5# (30) | NA | 4.8 ± 0.8** (24) |

| | **Mean distance from the OFM** (nm ± SD)[ab] (nb of measurements) | | | | |
|---|---|---|---|---|---|
| **Strain** | **CL** | **LM** | **DM** | **BLL** | **DSL** |
| *ΔspoIVB* | NA | NA | 31.3 ± 2.4 (24) | 55.1 ± 2.7 (24) | 81.3 ± 1.4 (24) |
| *ΔcotE*[c] | NA | NA | 27.4 ± 2.7** (30) | NA | 50.2 ± 2.2** (24) |

CW, cell wall; CL, crenelated layer; LM, light matrix; DM, dark matrix, BLL, bead-like layer; DSL, dark smooth layer; OFM, outer forespore membrane; NA, not applicable.

[a] Values in nm are presented as mean values ± standard deviations (with numbers of measurements in parentheses).

[b] Measured from the outer face the cytoplasmic membrane to the inner face of the given coat layer.

[c] *t* test was performed between corresponding coat layers in *ΔspoIVB* and *ΔcotE* sporangia. #, no statistical difference; *, $P < 0.05$; **, $P < 0.01$.

**Table S3. Dimensions of cell wall and coat layers measured from TEM of resin sections.**

| | **Mean thickness** (nm ± SD)[a] (nb of measurements) | | | | | | |
|---|---|---|---|---|---|---|---|
| **Strain** | **CW** | **Thin LM** | **Thin DM** | **Thick LM** | **Thick DM** | **Thin SL** | **Thick SL** |
| *ΔspoIVB* | 24.5 ± 2.4 (11) | 3.2 ± 0.2 (6) | 8.9 ± 0.8 (6) | 17.8 ± 3.7 (6) | 22.5 ± 4.5 (24) | 6.0 ± 1.0 (18) | 5.2 ± 1.0 (13) |
| *ΔcotE* [c] | 25.4 ± 1.9 (18)$^{\#}$ | 3.7 ± 0.5 (12)$^{*}$ | 5.1 ± 0.6 (12)$^{**}$ | 20.4 ± 3.9 (18)$^{\#}$ | 20.4 ± 2.2 (18)$^{\#}$ | NA | 4.7 ± 0.3 (12)$^{\#}$ |
| *ΔspoVID* [d] | 24.2 ± 3.4 (12)$^{\#}$ | NA | NA | 16.9 ± 2.3 (30)$^{\#}$ | 20.8 ± 4.0 (30)$^{\#}$ | 5.3 ± 0.9 (18)$^{*}$ | 4.9 ± 0.6 (6)$^{\#}$ |
| *ΔsafA* [e] | 23.1 ± 2.7 (18)$^{\#}$ | 3.5 ± 0.6 (12)$^{\#}$ | 5.3 ± 1.3 (12)$^{**}$ | NA | NA | 5.0 ± 0.5 (6)$^{\#}$ | 5.9 ± 1.1 (12)$^{\#}$ |

| | **Mean distance from the OFM** (nm ± SD)[ab] (nb of measurements) | | | | | |
|---|---|---|---|---|---|---|
| **Strain** | **Thin LM** | **Thin DM** | **Thick LM** | **Thick DM** | **Thin SL** | **Thick SL** |
| *ΔspoIVB* | NA | 3.2 ± 0.2 (6) | 12.1 ± 0.5 (6) | 31.0 ± 3.8 (24) | 56.3 ± 6.3 (13) | 60.4 ± 2.8 (12) |
| *ΔcotE* [c] | NA | 3.7 ± 0.5 (12)$^{\#}$ | 7.8 ± 1.2 (12)$^{\#}$ | 20.4 ± 3.9 (12)$^{*}$ | NA | 56.6 ± 1.5 (12)$^{**}$ |
| *ΔspoVID* [d] | NA | NA | NA | 17.1 ± 2.3 (24)$^{**}$ | 45.8 ± 2.6 (12)$^{**}$ | 71.3 ± 3.7 (12)$^{\#}$ |
| *ΔsafA* [e] | NA | 3.5 ± 0.6 (12)$^{\#}$ | 8.8 ± 0.9 (12)$^{**}$ | NA | NA | NA |

CW, cell wall; LM, light matrix; DM, dark matrix, SL, structured layer; OFM, outer forespore membrane; NA, not applicable.

[a] Values in nm are presented as mean values ± standard deviations (with numbers of measurements in parentheses).

[b] Measured from the outer face the OFM to the inner face of the given coat layer. For *ΔspoVID*, we excluded the arch-like structures of the thick dark matrix in our measurements.

[c, d ,e] *t* test was performed between corresponding coat layers in *ΔspoIVB* and *ΔcotE* ([c]), *ΔspoVID* ([d]) or *ΔsafA* ([e]) sporangia. #, no statistical difference; *, $P < 0.05$; **, $P < 0.01$.

#### SUPPORTING INFORMATION MOVIES

**Movie S1.** Sequential slices through a representative cryo-electron tomogram of a *ΔspoIVB B. subtilis* sporulating cell in cross-section view. In this stage-III sporangium, early stages of coat assembly are visible at the surface of the forespore. Related to Figure 1.

**Movie S2.** Sequential slices through a representative cryo-electron tomogram of a *ΔcotE B. subtilis* sporulating cell in cross-section view. In this stage-III sporangium, the DNA harbors a fibrillary toroid structure in the forespore cytoplasm. Related to Figure 2.

**Movie S3.** 3D rendering model built from the segmentation of the cryo-electron tomogram shown in Movie S1. Related to Figure 1.

**Movie S4.** 3D rendering model built from the segmentation of the DNA from the cryo-electron tomogram shown in Movie S2. Related to Figure 2.
